# Analgesic preference and injury behaviour in the cockroach *Blaptica dubia*

**DOI:** 10.64898/2026.06.02.729522

**Authors:** Rebecca D. Farquhar, David N. Fisher

## Abstract

Insects are increasingly recognised as behaviourally complex animals, yet whether they experience pain as an affective state beyond nociception remains unresolved. Voluntary ingestion of analgesics by injured individuals has been proposed as a key but largely untested criterion for evaluating insect pain. This study tested whether wing-clipped *Blaptica dubia* preferentially consumed an ibuprofen–sucrose solution over sucrose alone, and whether injury altered the duration of abnormal behaviour.. Adult males were assigned to injured or sham-handled groups and completed 30-minute two-choice assays.. Choice between the analgesic, sucrose-only and neutral zones were recorded, as well as recording the duration of a range of abnormal behaviours. The experiment was repeated with vanilla scent added to both solutions as a masking agent.. Injured cockroaches did not show greater analgesic preference in either condition. Instead, the vanilla condition markedly reduced preference for the analgesic.. Injury strongly increased the duration of abnormal behaviours, with increases in the duration of body flexion, lateral swaying, and wing fluttering, while wing-directed grooming was rare but showed a tendency for increases when injured. These findings provide no evidence that injured *B. dubia* actively seek an orally delivered analgesic under the conditions tested. However, injury produced clear and sustained behavioural disruption consistent with an internally driven aversive or discomfort-related state, highlighting both the methodological challenges of adapting voluntary analgesic assays to insects and the welfare relevance of injury in *B. dubia*.

## Introduction

Whether insects are capable of experiencing pain – defined as an affective state distinct from nociception^1^ – has become a central question in modern neuroethology and invertebrate welfare science^2,3^. While nociception refers specifically to the detection and neural processing of noxious stimuli and can occur without any accompanying affective experience, pain requires aa subjective unpleasant sensory and emotional experience^1^. Historically, insects were assumed to lack this capacity due to the comparatively limited size and organisational complexity of their brain and nervous systems, the presumed absence of integrative neural structures, and the lack of behavioural indicators consistent with negative affective states^4^ (but see below).

In animals inferred to experience pain, injury produces sustained changes in motivation, behaviour and decision-making that extend beyond immediate reflexive responses^5^. Therefore, nociceptive behaviours such as withdrawal reflexes cannot alone demonstrate pain. Instead, converging behavioural and physiological criteria, including central processing of noxious stimuli, learnt associations due to injury, and responsiveness to analgesics, are used to infer pain^5,6^. While no single criterion is sufficient on its own to demonstrate pain, advances in insect neurobiology and behaviour have challenged long-held assumptions by accumulating evidence from a range of sources. For example, insects possess specialised nociceptors responsive to harmful stimuli^7,8^ and injury induces sustained, non-reflexive behaviours such as body flexion, abdominal contractions and wound-directed grooming that persist beyond the initial stimulus^2,9^. Among these criteria, voluntary analgesic ingestion has been proposed as one of the strongest behavioural indicators of pain^10^, because only an organism experiencing a continued unpleasant internal state would be expected to seek relief. In vertebrates, uninjured individuals typically avoid analgesic-laced food or water because of aversive taste or side effects, whereas injured individuals selectively consume it, indicating that relief from discomfort outweighs these costs^10^). Equivalent studies in insects, however, remain extremely limited^2^.

Research of pharmacological effects on behaviour support the possibility of analgesia preference in insects. In the cockroach *Periplaneta americana*, morphine suppresses wound grooming and naloxone reverses this effect, suggesting the presence of opioid-sensitive pathways^9^. Similar responses occur in crustaceans and other insects^11^, while local anaesthetics reduce nocifensive responses across multiple invertebrate taxa^2^. However, in a self-administration study, honeybees (*Apis mellifera*) increased their consumption of both morphine and a pure sucrose solution following noxious stimulation, suggesting no preference for an analgesic but perhaps increased energy requirements following wounding^12^. Therefore, we are still lacking studies demonstrating voluntary analgesic self-administration, where injury-contingent preference rather than overall consumption is the critical measure.

Such studies are challenging as insect feeding behaviour is highly sensitive to gustatory and olfactory cues, meaning that bitterness, scent or unfamiliar additives can suppress feeding regardless of internal state^2,3^, limiting the utility of traditional preference paradigms. For this reason, most insect studies to date rely on forced administration of analgesics via injection or topical application, which introduces confounds such as sedation or motor impairment and may disrupt normal behaviour^11^. Therefore, the ideal study would examine voluntary ingestion of analgesics while controlling for the taste and scent of these compounds.

The present study aimed to address this gap by testing whether injured *Blaptica dubia* cockroaches show a measurable preference for ibuprofen-laced sucrose compared with uninjured controls, while explicitly assessing the role of flavour as a potential confounding factor. Ibuprofen was selected because it has documented physiological effects in terrestrial insects^13^ and is sufficiently water-soluble to be administered orally. Cockroaches provide an ideal system to explore whether insects engage in voluntary analgesic ingestion following injury as their nociceptive pathways respond to both opioids and non-steroidal anti-inflammatory drugs (NSAIDs) such as ibuprofen^9^, and their feeding preferences can be readily quantified in choice-based paradigms. However, no published study has yet examined whether cockroaches voluntarily ingest analgesics following injury, despite repeated calls to address this evidence gap^2,3^.

Two core aims guided the study:

- Determine whether injury influences preference for an ibuprofen–sucrose solution.
- Characterise the duration and composition of abnormal behaviours following injury.

From these aims, the following hypotheses were derived:

1. Injured cockroaches would show higher preference for ibuprofen-laced sucrose than uninjured individuals, which were expected to show no systematic preference.
2. Injured cockroaches would show greater analgesic preference when vanilla scent was added to the solutions as flavour masking should reduce aversive sensory effects associated with the analgesic solution.
3. Injured individuals would spend a greater proportion of time performing abnormal behaviours than uninjured individuals.

## Materials & Methods

### Subjects and Housing

Adult male *Blaptica dubia* from laboratory colonies maintained at the University of Aberdeen (UK) were used for all experiments. Colonies were maintained at a mean temperature of 28.9°C (range: 28–29°C) and a mean relative humidity of 36.1% (range: 27–45%) in a temperature- and humidity-controlled room, and housed in 48L polypropylene storage containers (610 × 400 × 315 mm, Solent Plastics, Romsey, UK), lined with egg cartons to provide shelter and ventilation. Individuals were supplied with dry dog kibble, carrot and water ad libitum. See Fisher^14^ for further details on stock maintenance. Only adult males were used to minimise variability related to sex or developmental stage and because their wings (which are greatly reduced in adult females) were required for the injury treatment.

Although invertebrates are not regulated under UK animal research legislation, all procedures were conducted in accordance with institutional guidelines for the care and use of animals at the University of Aberdeen and following the Association for the Study of Animal Behaviour^15^guidelines for the ethical treatment of nonhuman animals in behavioural research and teaching.

### Fasting and Pre-trial Handling

Groups of males (up to 13 at a time, depending on how many individuals were required per trial) were transferred into 0.9L polypropylene boxes (220 × 100 × 70 mm, Solent Plastics, Romsey, UK) and fasted for approximately 24 hours before testing to increase feeding motivation, with each container containing a small piece of egg carton for shelter. After fasting, individuals were moved into 0.3L polypropylene boxes (120 × 85 × 65 mm, Solent Plastics, Romsey, UK) for testing. Each 0.3L box held two cotton balls (∼25 mm diameter), positioned in opposite corners (Fig. 1) and saturated with either:

- sucrose-only solution (S)
- sucrose + ibuprofen solution (A)

Cotton balls were prepared prior to cockroach handling to minimise stress and were kept covered in small plastic dishes (77 × 44 mm) until placement inside testing containers to reduce evaporation.

### Solution Preparation

Ibuprofen was sourced from liquid capsules (200mg per capsule). Each capsule was diluted in 10mL distilled water to produce a 20mg/mL stock. Trial solutions were mixed fresh in 500mL volumes:

- **Analgesic (A):** 500mL distilled water + 50mg sucrose + 200mg ibuprofen
- **Control (S):** 500mL distilled water + 50mg sucrose
- **Vanilla trials:** 0.25mL vanilla oil added to each A and S 500mL solution (0.05% v/v) to mask bitterness across treatments.

This produced a final ibuprofen concentration of 0.4mg/mL in the analgesic solution. Because no validated oral ibuprofen dose has been established for *B.dubia*, 0.4mg/mL was selected as a conservative exploratory concentration for use in a voluntary feeding assay. The concentration was kept relatively low to minimise risk of adverse effects or strong gustatory aversion suppressing consumption independently of injury. This decision was also informed by preliminary observations in which some individuals approached and contacted the analgesic cotton ball but withdrew almost immediately without sustained feeding, suggesting that the solution was detectable and may have had aversive sensory properties.

**Figure 1.**
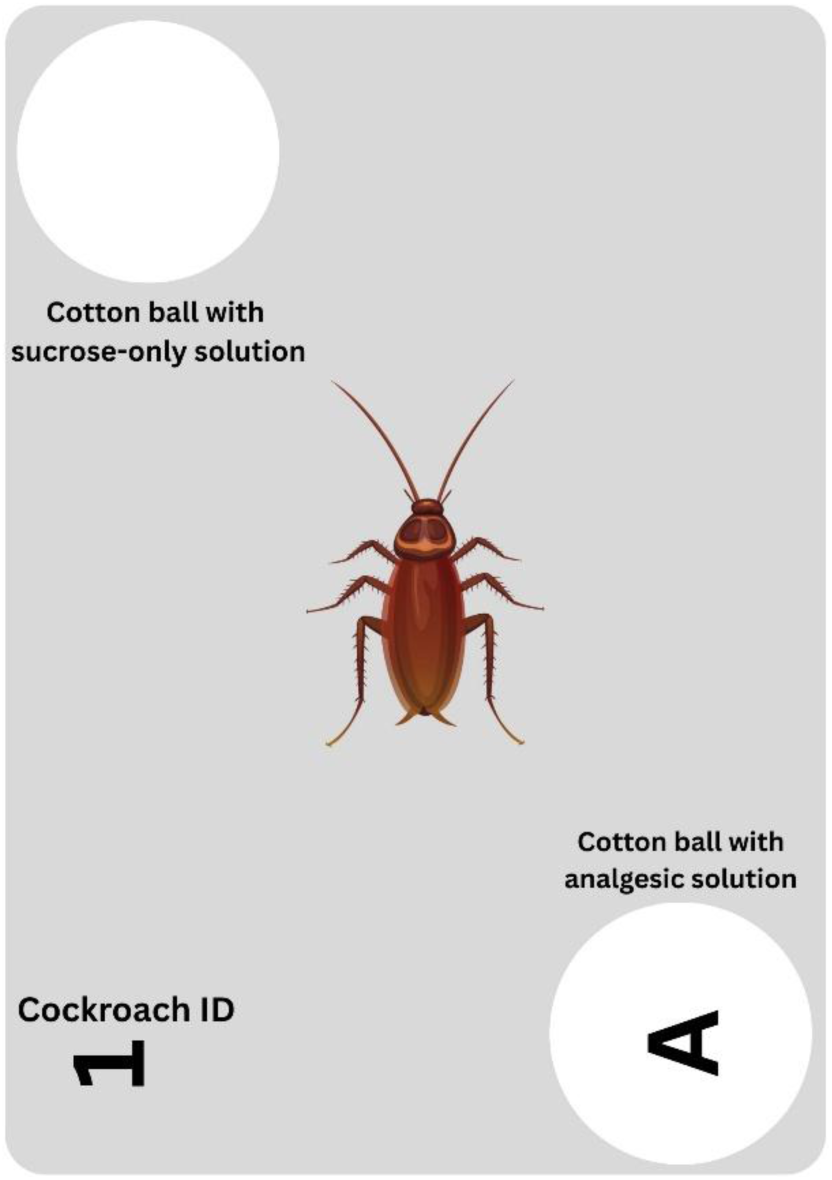
Diagram of the behavioural arena used during 30 min feeding-choice trials. Each 0.3L container contained two cotton balls positioned in opposite corners: one saturated with sucrose-only solution (control) and one with ibuprofen–sucrose solution (labelled “A “for analgesic).

Fifteen millilitres of each solution were applied to the appropriate cotton ball immediately before each trial. Cotton balls were fully saturated to counteract evaporation; however, this occasionally resulted in solution spreading slightly beyond the perimeter of the cotton ball within the trial arena.

### Injury Procedure & Handling

A mild injury was induced using sterilised dissection scissors. Approximately 10mm of the tip of the male cockroach’s wings was removed, following the approach of Jenkinson *et al*.^16^. This produced a consistent injury without impairing locomotion.

Uninjured cockroaches were put through sham procedures with the same handling method and with contact from scissors to wing, but without their wings being clipped, to avoid introducing differences caused by handling or stress rather than the injury itself.

Standard PPE (lab coat and gloves) was worn throughout. Ethical handling procedures were followed at all times, including working over a 48L box to prevent escape or accidental injury if an individual was dropped, handling animals gently but securely and minimising handling time to reduce stress.

### Experimental Design

Trials were conducted across two experimental conditions: a default, control condition and a separate vanilla-flavoured condition, in which 0.25mL vanilla oil was added to both the analgesic and sucrose-only solutions to mask bitterness. Trials consisted of mixed groups of injured and uninjured males, with numbers ranging from 8 individuals (e.g. 4 injured + 4 uninjured) early in the experiment while the workflow was being refined, to a maximum of 26 individuals (13 injured + 13 uninjured) towards the end of the experiment. In total 50 individuals were tested in each of the plain and vanilla-scented conditions.

Each cockroach was tested individually in a 0.3L box placed inside a larger 48L box on top of a pet heating mat (models TK-HPP7040A & TK-HPP6540, OnKey Electronic Technology Co. Ltd, Dongguan, China) to reduce external disturbance and maintain consistent lighting and thermal conditions. Up to nine 0.3L boxes were placed in each 48L box, with up to three 48L boxes used per trial.

### Recording Conditions

Trials were recorded using IP 2MPx Wi-Fi mini tube cameras (ABUS, Wetter, Germany). Video was captured using the AnyCam IP camera software at 720p resolution (1280 × 720, anycam.io) and 25fps. Between one and three cameras were used per trial, one per 48L box being used. Cameras were mounted directly above each 48L box to provide an unobstructed overhead view of all 0.3L individual boxes within.

Before each trial, both the camera-room temperature and the internal temperature of the 48L boxes were measured using digital rod thermometers. Across trial days, the recording room was maintained at a mean temperature of 24.8°C (range: 24–25°C) and a mean relative humidity of 27.1% (range: 20–30%), and the 48L trial boxes were maintained at a mean temperature of 28.2°C (range: 27–29°C). Once cameras were aligned, room lights were switched off to encourage more natural behaviour in the cockroaches, which are primarily nocturnal^17^, and recordings were conducted in infrared mode to ensure consistent illumination. Each trial ran for 30 minutes, with additional 30 seconds before and after the trial period to ensure a clean and uninterrupted behavioural window. Because individuals had no prior exposure to the analgesic solution before testing, any association between its taste and potential analgesic effects would need to occur within the 30-minute trial period, a point we return to in the Discussion.

### Cleaning Procedures

All boxes, plastic dishes, and tools were cleaned with 70% ethanol after each use. Dishes used for solution application were additionally washed with dish soap to remove sucrose residue and prevent contamination.

### Post-trial Housing and Euthanasia

Uninjured individuals were returned to colony housing to maintain breeding stock. Injured individuals were euthanised by freezing at -20°C inside a sealed biowaste bag in accordance with institutional ethical and technician guidelines. Although the initial plan was to euthanise all individuals to avoid re-exposure, limited colony stock required retention of uninjured individuals. This means that some cockroaches may have been cryptically re-used across trials. Trials were conducted intermittently across the week, typically following an overnight fasting period, with no testing performed during weekends. Consequently, repeated exposure was generally separated by several days, although in some cases re-use may have occurred within approximately 1-2 days. Because cockroaches are capable of forming associative memories that can persist for at least 24 hours to several days or longer (depending on the conditioning paradigm and species^18,19^), prior exposure to handling, the apparatus, or the test solutions may have influenced subsequent behaviour.

### Behavioural Scoring

Behavioural videos were analysed manually using VLC Media Player videolan.org). Behaviour was scored at 30-second intervals, producing 61 observations per individual. At each timepoint, individuals were categorised as belonging to one of three zones:

- **A:** positioned at the analgesic cotton ball
- **S:** positioned at the sucrose-only cotton ball
- **N:** located within the neutral zone

Because the cotton balls were fully saturated and solution occasionally spread beyond the cotton surface, an individual was classified as interacting with A or S only when it was:

- oriented toward the cotton ball, and
- positioned with its head within ∼1cm of the cotton ball perimeter.

Individuals that were close to a cotton ball but facing away from it were scored as neutral. Neutral observations were retained because they allowed quantification of time spent away from both food sources rather than assuming not being near one meant showing a preference for the other.

In addition to spatial position (A, S or N), behaviours indicative of injury-related disruption were recorded. These behaviours were selected based on previous work showing that injury in insects can induce sustained, non-reflexive responses such as body flexion, abdominal contractions and wound-directed grooming^2,9^. Recording these behaviours allowed assessment of injury effects that occurred independently of feeding behaviour. These behaviours were extracted from the video in a separate pass to that used to determine the locations used for the preference assessment.

This enabled us to record the onset and cessation of each behavioural episode, giving a precise duration rather than estimating behaviour every 30s. For each individual, we recorded durations of body flexion (BF), lateral abnormal swaying (LAS), wing-directed grooming (WDG), wing fluttering (WF), and inversion (I).

Definitions and other notes for each of these are given in Table 1. Individuals that did not display any of a particular behaviour were assigned a duration of zero for that behaviour.

### Statistical Analysis

All analyses were conducted in R ver. 4.4.2^20^. Statistical analyses were designed to test for effects of experimental condition (Control vs. Vanilla) and injury status (Injured vs. Uninjured) on feeding behaviour and abnormal behavioural responses.

**Table 1.**
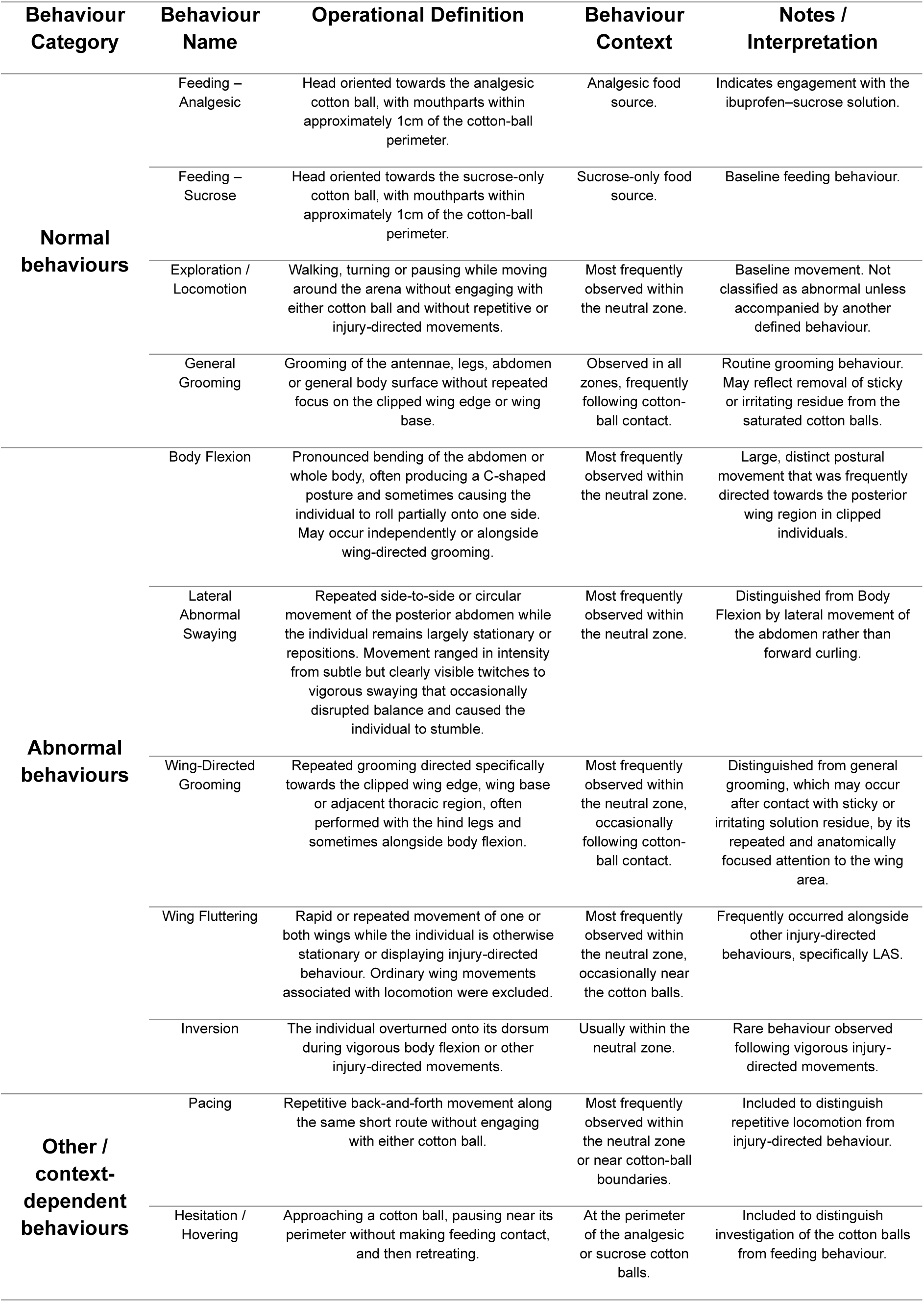
Ethogram of behaviours recorded during feeding-choice trials, giving operational definitions, contextual descriptions, and other notes on behaviours observed in *Blaptica dubia* during behavioural scoring. Behaviours were categorised as normal behaviours, abnormal behaviours associated with injury, and other behaviours.

### Preference Index Analysis

Analgesic preference was quantified using a preference index calculated as:

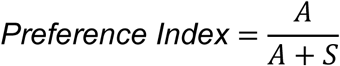

where A and S represent the number of observations at the analgesic and sucrose-only cotton balls respectively. Individuals with zero A and zero S counts (i.e., those that did not visit either cotton ball) were excluded from this analysis (n = 1 for Control-Uninjured, n = 3 for Control-Injured, n = 4 for Vanilla-Uninjured, and n = 2 for Vanilla Injured).

A Kruskal-Wallis test was used to determine whether preference indices differed across the four experimental groups (all combinations of experiment × injury).). Once confirming this test was significant, we followed up with pairwise Wilcoxon tests in the package rstatix^21^, with a Holm correction for multiple testing tests (giving *p* adjusted values [*pa*]). Four one-sample Wilcoxon signed-rank tests were also used to determine whether preference index within each of the four experiment × injury groups differed from 0.5, representing no.. Due to multiple testing the p values of four tests were adjusted using the Holm method.

### Abnormal Behaviour Measures

All 100 individuals were included in the abnormal behaviour analyses. Total abnormal behaviour duration was calculated by summing the duration of all scored abnormal behaviour types performed by each individual during the 30-minute trial. Individuals that did not display any abnormal behaviour received a duration of 0s.

Differences among the four experiment × injury groups for total abnormal behaviour duration and the duration of each of the different behaviours were assessed using Kruskal–Wallis rank-sum tests as for the preference index. Where overall differences were significant, pairwise Wilcoxon rank-sum tests with Holm correction were used. Note the frequency of inversion (only a single individual showed this quite extreme behaviour, see Fig. 2) precluded its analysis, while for wing-directed grooming no uninjured cockroaches displaying this behaviour and so all pairwise comparisons could not be performed. Instead, for wing-directed grooming following the Kruskal-Wallis test we only compared injured and uninjured in each of the experiments with Wilcoxon tests.

All analyses used a significance threshold of α=0.05. All figures were generated using ggplot2^22^.

**Figure 2.**
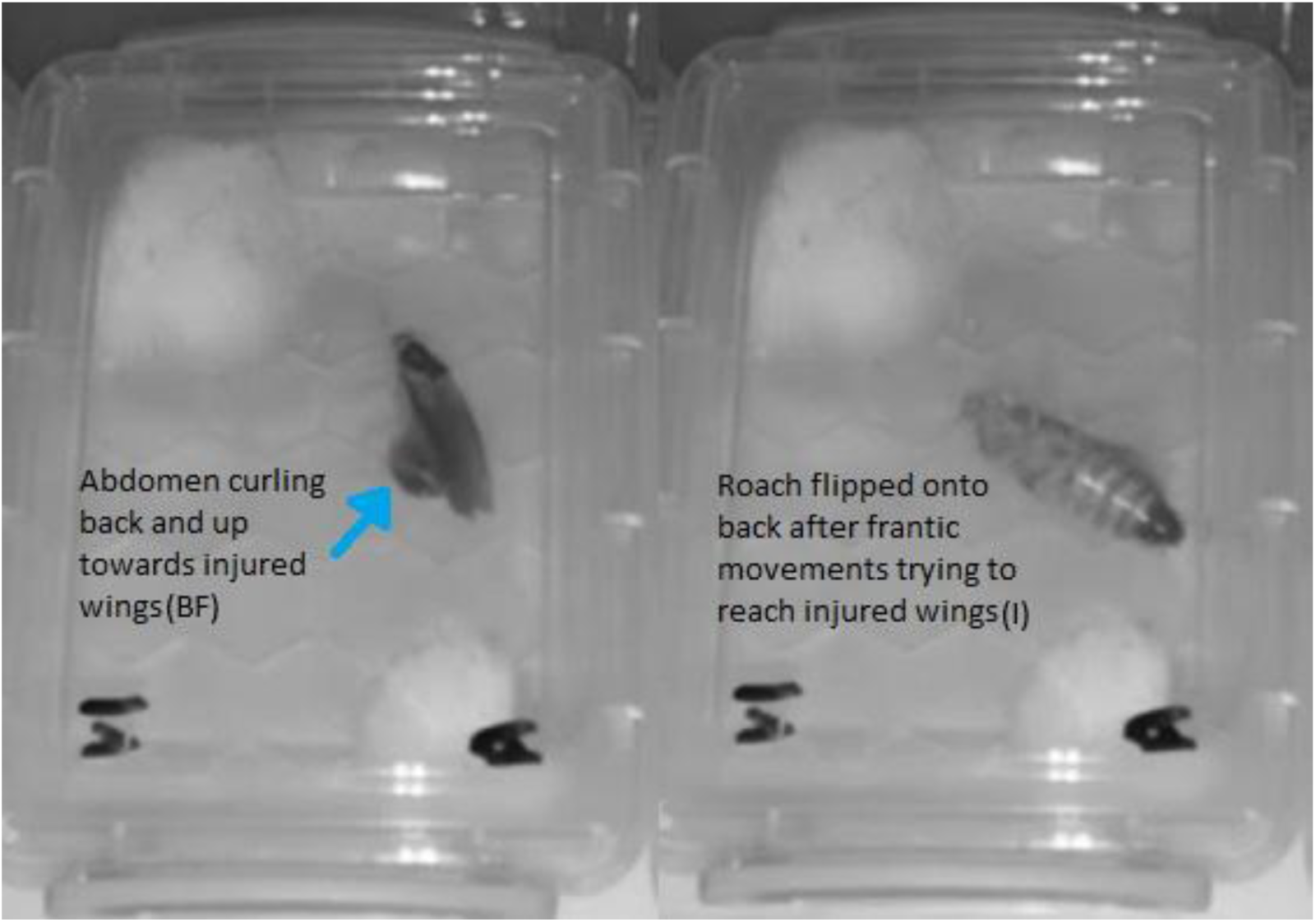
Inversion observed in a single Vanilla Injured cockroach (V1) during injury-directed behaviour. Sequential frames show the individual overturning onto its dorsum (“inversion”) following vigorous body flexion and contortion directed towards the clipped wing. The inversion lasted approximately 11s before the individual righted itself.

## Results

### Results summary

In both experiments injury status had little effect on analgesic preference, but while control cockroaches preferred the analgesic solution over sucrose only, the vanilla cockroaches showed the opposite preference. In contrast, the duration of abnormal behaviours was strongly associated with injury, with injured cockroaches displaying more persistent and frequent abnormal behaviours than uninjured individuals, with presence of the vanilla having little effect.

### Preference for analgesia

Preference indices differed among experimental groups (Kruskal-Wallis test, χ²(3)=53.02, *p*<.001), being uniformly higher in the control experiment than in the vanilla experiment (Fig. 3, Table 2). The only pairs not different were Control-Injured vs. Control-Uninjured, and Vanilla-Injured vs. Vanilla-Uninjured (Table 2). Preferences in the control experiment were above 0.5 for both uninjured (median=0.730, Wilcoxon test, W=252, *p_a_*=.001) and injured individuals (median=0.690, Wilcoxon test, W=188, *p_a_*=.012). Preferences in the vanilla experiment were below 0.5 for both uninjured (median=0.071, Wilcoxon test, W=7, *p_a_*<.001) and injured individuals (median = 0.125, Wilcoxon test, W=1, *p_a_*<.001).

### Abnormal behaviour

Abnormal behaviour was observed in 80% of injured individuals compared with 34% of uninjured individuals (6/25 Control Uninjured, 19/25 Control Injured, 11/25 Vanilla Uninjured and 21/25 Vanilla Injured cockroaches). Total abnormal behaviour duration differed significantly among the four experimental groups (Kruskal–Wallis test: χ²(3)=34.98, *p*<.001; Figure 4). Injured individuals displayed substantially longer durations of abnormal behaviour than uninjured individuals, with median abnormal behaviour duration being 0s in the Control uninjured group, 139s in the Control **g** group, 0s in the Vanilla Uninjured group and 233s in the Vanilla Injured group (full pairwise comparisons in Table 2).

BF and LAS accounted for the majority of abnormal behaviour duration in injured cockroaches, whereas WF and WDG contributed less to total duration (Figure 5). BF occurred in 30/50 injured and 12/50 uninjured cockroaches, while LAS occurred in 35/50 and 7/50 uninjured cockroaches. WDG was observed in 8/50 injured cockroaches and was not observed in uninjured cockroaches. WF occurred in 25/50 injured and 3/50 uninjured cockroaches.

**Figure 3.**
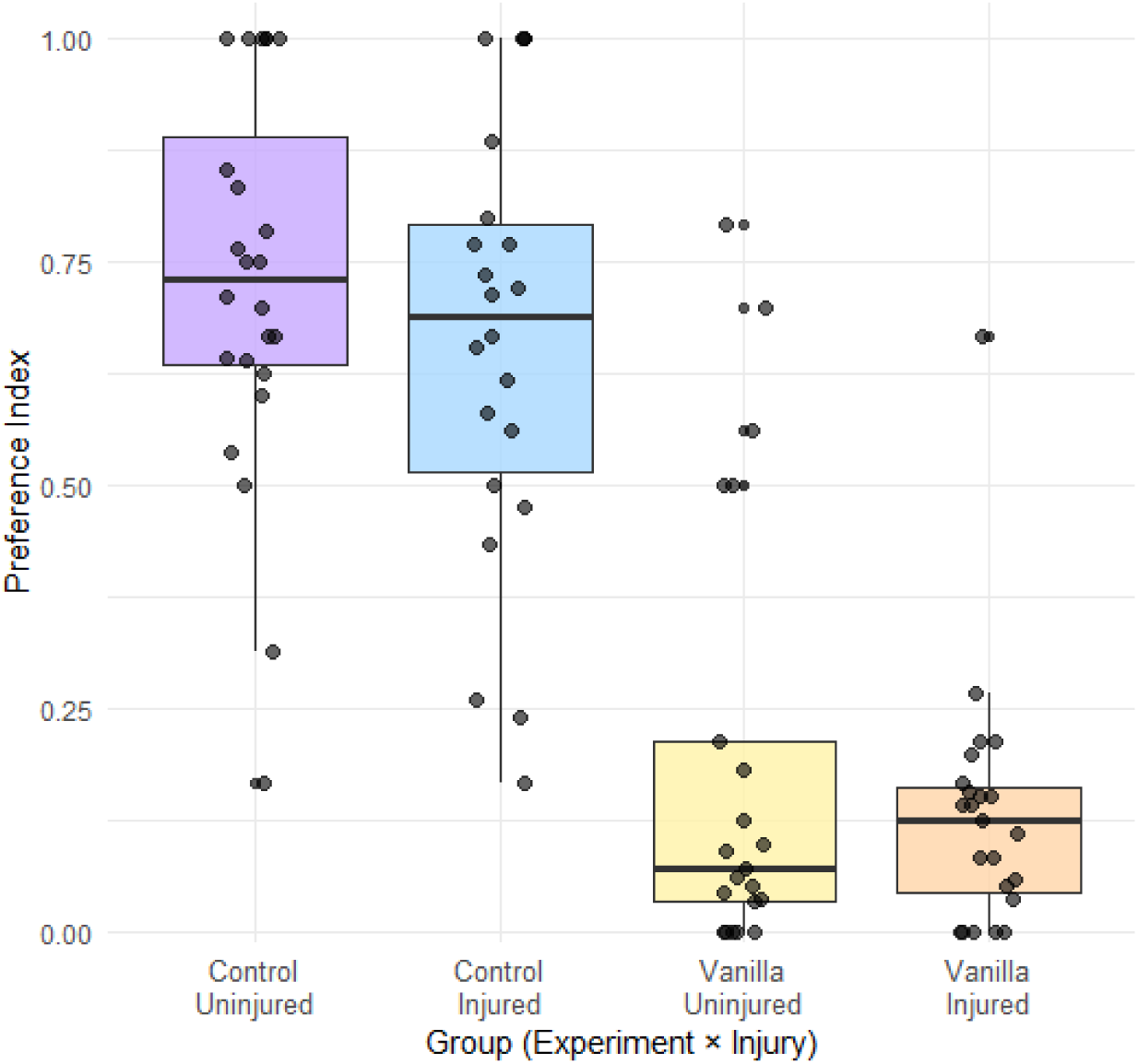
Preference index (count of visits to analgesic solution / count of visits to either solution) across the four experimental groups (Control Uninjured, Control Injured, Vanilla Uninjured, and Vanilla Injured). Boxplots show the median, interquartile range, and distribution. Control cockroaches showed higher preference index values than vanilla cockroaches, with injury status having minimal effect.

BF duration differed among experimental groups (Kruskal-Wallis test, χ^2^(3)=13.806, *p*=.003; Figure 5a), being higher in both injured groups compared to the control uninjured group. BF did not differ among the other pairs (Table 2). LAS differed between groups (Kruskal-Wallis test, χ^2^(3)=37.245, *p*<.001; Figure 5b) based on injury, being higher in both injured groups than both uninjured groups, but did not differ based on experimental context (Table 2). WF was different among experimental groups (Kruskal-Wallis test, χ^2^(3)=22.765, *p*<.001; Figure 5c), being higher in both injured groups than both uninjured groups, but not differing between the two injured groups or the two uninjured groups (Table 2). Finally, WDG was different among groups (Kruskal-Wallis test, χ^2^(3)=8.593, *p=*.035; Figure 5d), being totally absent in both uninjured groups. Each injured group performed more WDG than the uninjured group in the same experiment (Table 2).

**Table 2.**
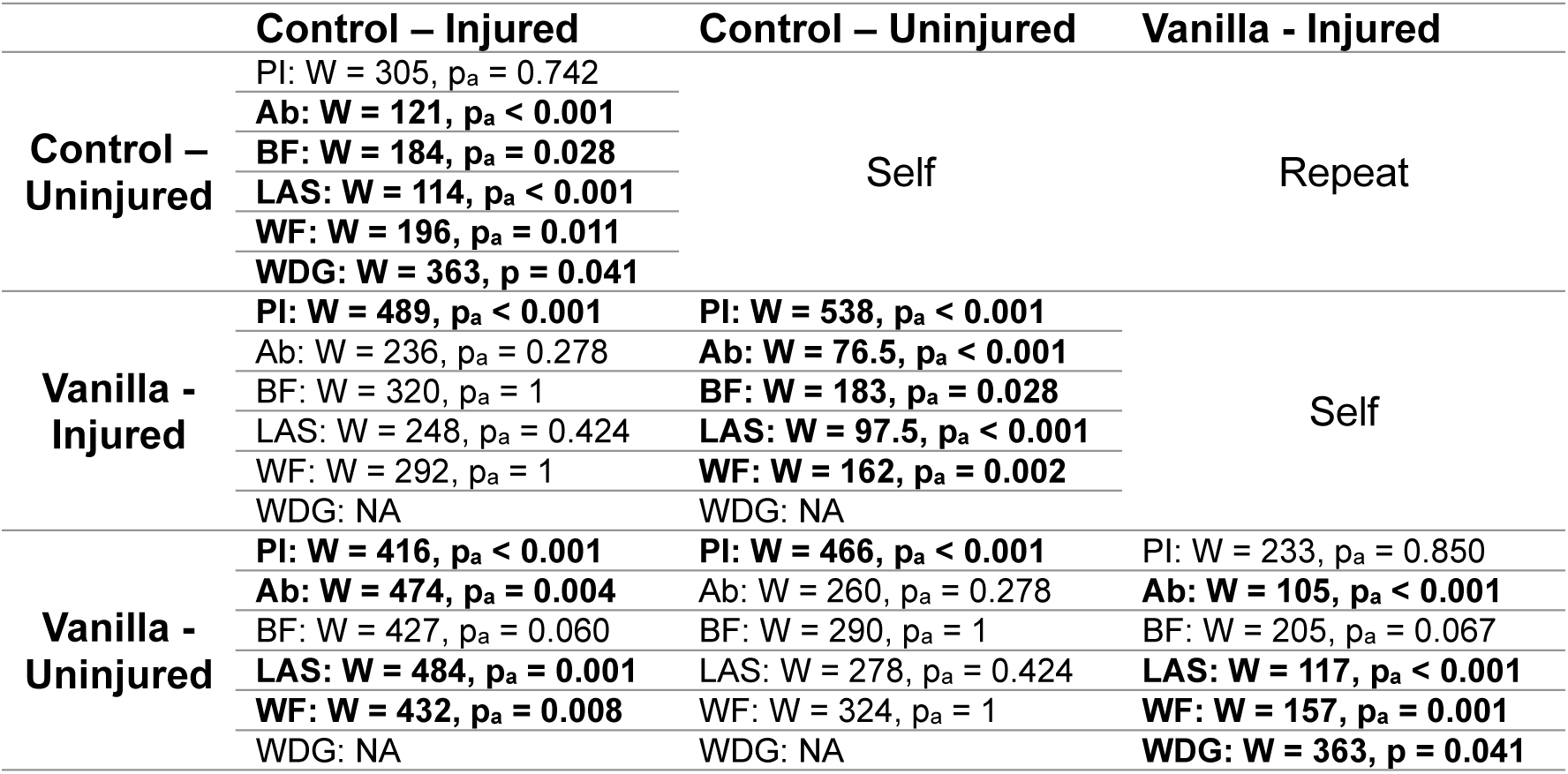
A matrix presenting the results of the Wilcoxon pairwise comparisons among experimental groups for the preference index (PI), total abnormal behaviour duration (Ab), duration of body flexion (BF), duration of lateral abnormal swaying (LAS), duration of wing fluttering (WF), and duration of wing-directed grooming (WDG). The large cells show the intersection between the four different experiment groups (across cells and down rows, 3out of 4 shown to remove redundancy) and within the cells show the test statistic and p value of the Wilcoxon comparison for each behaviour. For sample sizes in each comparison see main text. For all comparisons except for WDG (which had too few non-zero values to perform all comparisons) we give “p_a_” values, which are adjusted p values after a Holm correction for multiple testing. Statistically significant comparisons (p or p_a_ < 0.05) are highlighted in bold.

|  | Control – Injured | Control – Uninjured | Vanilla - Injured |
| --- | --- | --- | --- |
| **Control – Uninjured** | PI: W = 305, $p_a = 0.742$ | Self | Repeat |
|  | **Ab: W = 121, $p_a < 0.001$** |  |  |
|  | **BF: W = 184, $p_a = 0.028$** |  |  |
|  | **LAS: W = 114, $p_a < 0.001$** |  |  |
|  | **WF: W = 196, $p_a = 0.011$** |  |  |
|  | **WDG: W = 363, $p = 0.041$** |  |  |
| **Vanilla - Injured** | **PI: W = 489, $p_a < 0.001$** | **PI: W = 538, $p_a < 0.001$** | Self |
| | Ab: W = 236, $p_a = 0.278$ | **Ab: W = 76.5, $p_a < 0.001$** | |
| | BF: W = 320, $p_a = 1$ | **BF: W = 183, $p_a = 0.028$** | |
| | LAS: W = 248, $p_a = 0.424$ | **LAS: W = 97.5, $p_a < 0.001$** | |
| | WF: W = 292, $p_a = 1$ | **WF: W = 162, $p_a = 0.002$** | |
|  | WDG: NA | WDG: NA |  |
| **Vanilla - Uninjured** | **PI: W = 416, $p_a < 0.001$** | **PI: W = 466, $p_a < 0.001$** | PI: W = 233, $p_a = 0.850$ |
| | **Ab: W = 474, $p_a = 0.004$** | Ab: W = 260, $p_a = 0.278$ | **Ab: W = 105, $p_a < 0.001$** |
| | BF: W = 427, $p_a = 0.060$ | BF: W = 290, $p_a = 1$ | BF: W = 205, $p_a = 0.067$ |
| | **LAS: W = 484, $p_a = 0.001$** | LAS: W = 278, $p_a = 0.424$ | **LAS: W = 117, $p_a < 0.001$** |
| | **WF: W = 432, $p_a = 0.008$** | WF: W = 324, $p_a = 1$ | **WF: W = 157, $p_a = 0.001$** |
|  | WDG: NA | WDG: NA | **WDG: W = 363, $p = 0.041$** |

## Discussion

This study investigated whether injured *Blaptica dubia* would exhibit voluntary analgesic ingestion and whether injury would induce persistent abnormal behaviours independent of feeding. Injured cockroaches did not show a measurable preference for the ibuprofen-laced solution; instead, feeding behaviour was strongly influenced by experimental condition, with individuals in the control experiment generally preferring the analgesic solution, but individuals in the vanilla experiment experiment avoiding the analgesic in favour of just sucrose. In contrast, injury consistently increased the duration of abnormal behaviours, with BF and LAS accounting for most recorded abnormal behaviour. These findings indicate that, under the conditions tested, voluntary analgesic preference was not detected, whereas behavioural disruption following injury was robust and sustained.

**Figure 4.**
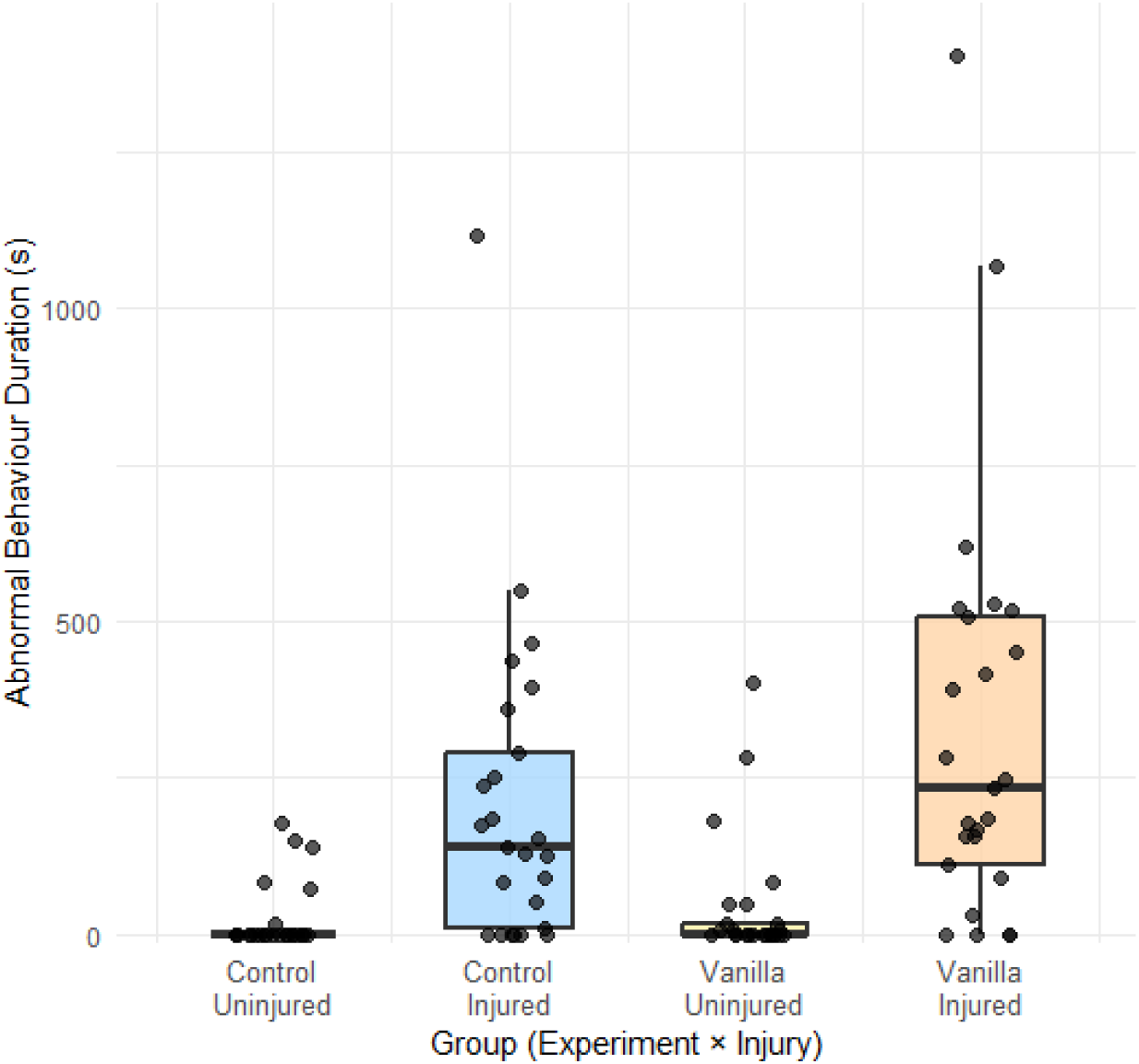
Total abnormal behaviour duration (seconds) across the four experiment × injury groups (Control Uninjured, Control Injured, Vanilla Uninjured, and Vanilla Injured). Boxplots show the median, interquartile range, and range (excluding outliers) with individual observations overlaid. Injured cockroaches displayed significantly longer durations of abnormal behaviours than uninjured cockroaches.

In the context of insect pain research, a central question is whether injured individuals voluntarily ingest pain-relieving solutions when offered a choice between these and control solutions. Although insects exhibit nociception, wound-directed grooming, learning in response to injury, and pharmacological responsiveness to analgesics^2,23^, voluntary analgesic self-administration – one of the strongest behavioural indicators of pain used in vertebrate research – has remained almost entirely untested in insects. By adapting this paradigm to *B. dubia*, we aimed to evaluate whether injured cockroaches would preferentially consume ibuprofen-laced sucrose to assess the feasibility of transferring vertebrate-style choice assays to an invertebrate context. In addition, we recorded abnormal behaviours to determine whether injury produced broader behavioural changes that could provide further evidence of pain.

**Figure 5.**
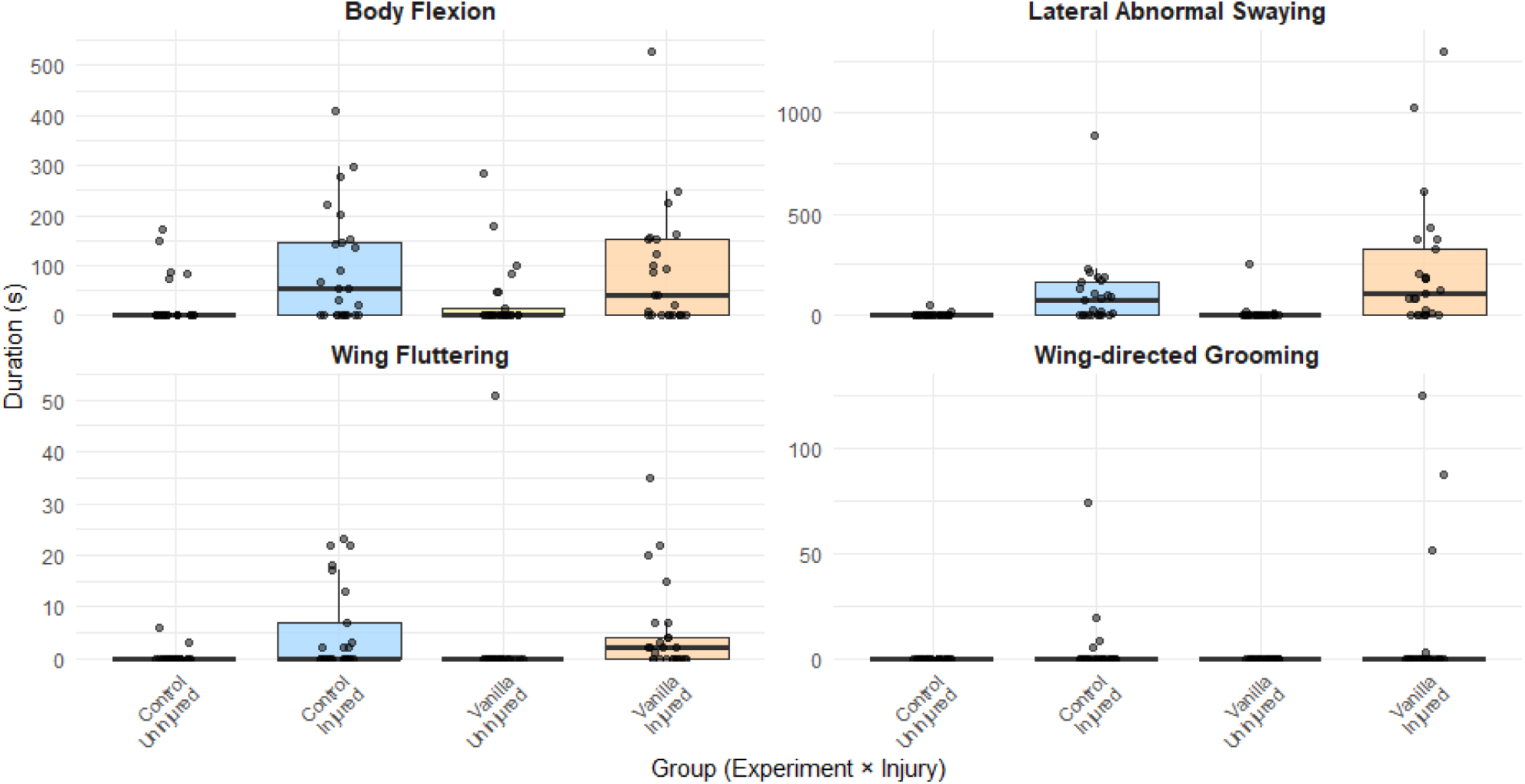
Duration of individual abnormal behaviour types (seconds) across the four experiment × injury groups (Control Uninjured, Control Injured, Vanilla Uninjured, and Vanilla Injured). Separate panels show body flexion, lateral abnormal swaying, wing fluttering, and wing-directed grooming. Boxplots show the median and interquartile range, with individual observations overlaid. Note that y-axis scales differ among panels.

Across both experimental batches, injured cockroaches did not demonstrate a measurable preference for the ibuprofen-laced solution. Instead, feeding behaviour was dominated by a scent effect: cockroaches in the control experiment preferred the analgesic solution regardless of injury, while those in the vanilla condition had the opposite preference. This reduction cannot be attributed to increased inactivity or neutral-zone occupancy, because the preference index was calculated using only visits to the analgesic and sucrose cotton balls. The vanilla additive, intended to improve palatability, appears instead to have interacted with the ibuprofen to suppress feeding in both injured and uninjured individuals. Ethological notes support this interpretation, as vanilla individuals often engaged in immediate grooming after contacting the cotton balls, suggesting that the additive rendered the solutions aversive. This pattern indicates that sensory interference – arising from altered palatability, texture or detectability – likely masked any subtle motivational differences linked to injury^2,3^.

These findings illustrate a fundamental challenge in applying vertebrate pain paradigms to insects. Insects adhere to strict palatability thresholds and can reject normally attractive food sources when gustatory processing is altered; for example, cockroaches can develop glucose aversion following changes in taste receptor function, causing sugars that are typically rewarding to become behaviourally aversive^24^. Without first establishing the taste detectability, solubility and behavioural neutrality of candidate analgesics, the absence of preference cannot be interpreted as a meaningful indicator of internal state. The present findings therefore highlight the importance of refining methodological aspects of voluntary analgesic assays in insects, as sensory properties of test compounds may substantially influence feeding behaviour and reduce the sensitivity of preference-based measures.

A further limitation is that the ibuprofen concentration used may have been too low to produce detectable analgesic effects in *B. dubia*. Cammaerts and Cammaerts^13^ reported behavioural and physiological effects in the ant *Myrmica sabuleti* following chronic exposure to a substantially higher ibuprofen concentration than the 0.4mg/mL used here. However, differences in species, exposure duration and experimental purpose prevent direct comparison between the two studies. Because ibuprofen had not previously been tested in *B. dubia*, a relatively low concentration was selected initially to minimise the risk of adverse effects or strong sensory aversion. Early pilot observations showed that some cockroaches approached the analgesic cotton ball, paused near its perimeter and then withdrew without feeding, suggesting that the solution may have been detectable or aversive. The concentration was therefore not increased further, and a subsequent experimental condition included vanilla as a masking agent. However, this cautious approach may have reduced the likelihood that individuals ingested a pharmacologically effective dose.

Another methodological consideration is the time required for animals to associate analgesic ingestion with relief from discomfort. In many vertebrate studies demonstrating analgesic self-administration, individuals experience the effects of the analgesic before the experimental choice tests begin, allowing them to learn that ingestion produces relief^10^. In the present study, the first opportunity for cockroaches to ingest ibuprofen occurred during the recorded trials themselves. Because NSAIDs such as ibuprofen may take time to produce physiological effects, the 30-minute observation window may not have allowed sufficient time for individuals to experience analgesic relief and subsequently alter their feeding behaviour. If cockroaches require repeated exposure to associate the taste of the analgesic solution with reduced discomfort, the experimental design may therefore have limited the likelihood of detecting injury-contingent preference. However, this limitation is unlikely to reflect an inability to form such associations, as cockroaches are capable of associative learning and can retain learned information ranging from several days to weeks^18,19^. Instead, the limited duration and single-exposure nature of the present assay may simply have provided insufficient opportunity for individuals to experience analgesic effects and subsequently modify their behaviour.

Because uninjured individuals were returned to the communal colony between trials and were not individually marked, it is possible that some cockroaches were inadvertently reused across trials. Cockroaches are capable of forming associative memories lasting from several days to weeks, depending on the training paradigm^18,19^. Although trials in this study were typically spaced a few days apart, the full extent of memory retention in *B. dubia* remains unknown. It is therefore plausible that prior handling, apparatus exposure, stress, or prior exposure to the test solutions – including the analgesic – may have influenced behaviour in subsequent trials. Such untracked individual histories introduce uncontrolled variation, limiting the interpretability of negative findings, rather than introducing any systematic bias.

Despite these complications, the data on abnormal behaviours provided robust insights. Injured individuals displayed prolonged abnormal behaviours – especially BF and LAS, but also WDG and WF – that differed markedly from the brief contamination-related grooming seen in uninjured cockroaches. These behaviours frequently dominated time spent away from feeding sites, suggesting a motivational conflict in which internally driven responses to injury overrode feeding or exploration. This pattern suggests is consistent with injury influencing behavioural priorities, resulting in reduced engagement with feeding-related activities. During the quantification of the durations of each abnormal behaviour, it became apparent that BF was often associated with WDG, suggesting that these movements may represent complementary responses directed towards the injured wing rather than entirely independent behaviours. Although these behaviours cannot be interpreted as unequivocal evidence of pain, they are consistent with a persistent state of discomfort or irritation following injury. The marked increase in their duration, together with their repeated direction towards the clipped wing, suggests that injury altered behavioural priorities independently of feeding.

The behaviour of individual V1 provides a particularly striking example (Fig. 2). Although inversion was observed only once, it illustrates the intensity that injury-directed behaviour could occasionally reach. In attempting to reach the clipped wing tip, V1 flexed and twisted its abdomen with sufficient force that it eventually overturned onto its dorsum. The inversion observed here was not deliberate but occurred as a consequence of a sustained cluster of intense BF and LAS, including several episodes lasting 35-51 seconds. A 20 second BF episode immediately preceded the 11 second inversion, after which a further prolonged episode of LAS occurred. This sequence suggests that the inversion was not an isolated loss of balance but occurred within a sustained series of vigorous movements associated with the clipped wing region. Although inversion was observed in only one individual and cannot itself demonstrate pain, the persistence and intensity of the surrounding behaviours indicate a substantial response to injury, potentially reflecting discomfort, irritation or altered sensation. The movements became sufficiently forceful to disrupt normal posture and mobility, suggesting that attention towards the injured area temporarily outweighed competing behavioural priorities. In a natural setting, overturning onto the dorsum could leave a cockroach highly vulnerable by restricting movement, delaying escape and increasing exposure to predators or other threats until it was able to right itself. The fact that V1 continued to perform intense injury-directed behaviour despite these apparent costs suggests that the response was strongly motivated. However, this single observation cannot be taken as evidence of pain by itself and may instead reflect intense discomfort, irritation or altered sensation associated with the injury. Similarly, although brief pauses sometimes followed contact with the ibuprofen cotton ball, these events cannot be interpreted as evidence of pharmacological relief, as they may simply represent a temporary interruption to the ongoing injury-directed behaviour. Together, these observations illustrate how feeding interactions and injury-associated behaviours could occur within the same trial without demonstrating analgesic-seeking behaviour.

Taken together, these behavioural observations are consistent with several indicators commonly used to infer pain-like states in invertebrates^2,5,6^. The responses were persistent, directed toward the site of injury and occurred repeatedly throughout the trial rather than appearing as brief reflexive reactions. Although the voluntary analgesic-ingestion assay did not demonstrate injury-contingent preference under the conditions used here, the abnormal behaviours observed suggest that injury produced a sustained internal disturbance that affected behavioural priorities.

These findings highlight both the promise and the methodological constraints of applying voluntary analgesic-choice paradigms to insects. Before such tests can reliably evaluate relief-seeking behaviour, future studies should establish the palatability and detectability of candidate analgesics and masking agents in insect feeding systems and consider experimental designs that allow individuals to experience analgesic effects before preference tests begin. Repeated exposure may be particularly important if individuals must learn to associate the sensory properties of an analgesic solution with subsequent relief. Improved tracking of individual animals and longer observation periods may also help clarify whether insects alter their feeding behaviour after experiencing pharmacological relief over longer timescales.

## Conclusion

Overall, although the present study did not provide evidence that injured *B. dubia* preferentially consumed an analgesic under the conditions tested, it did reveal clear and sustained behavioural disruption following injury. Injured cockroaches displayed significantly longer durations and more frequent episodes of abnormal behaviour, with BF and LAS accounting for most of the recorded response and WF and WDG occurring less frequently. These behaviours were persistent and often directed toward the injured wing region, suggesting a sustained state of discomfort, irritation, or another negatively valanced internal state, although they cannot be taken as direct evidence of pain without further research into the motivation underlying these behaviours. These findings illustrate both the potential and practical limitations of applying voluntary analgesic-ingestion paradigms to insects and contribute to ongoing investigations into insect pain and welfare.

## Acknowledgements

We thank Keith Lockhart for his extensive efforts maintaining the *Blaptica dubia* stock population.

## References

1. Raja, S.N. et al. (2020) ‘The revised International Association for the Study of Pain definition of pain: concepts, challenges, and compromises’, Pain, 161(9), pp. 1976–1982. Available at: 10.1097/j.pain.0000000000001939

2. Gibbons, M., Chittka, L. and Barron, A.B. (2022) ‘Can insects feel pain?’, Advances in Insect Physiology, 63, pp. 155–229. Available at: 10.1016/bs.aiip.2022.10.001

3. Crump, A., Browning, H., Schnell, A.K., Burn, C. and Birch, J. (2022) ‘Sentience in decapod crustaceans: A general framework and review of the evidence’, Animal Sentience, 32(1), pp. 1–65. Available at: https://www.wellbeingintlstudiesrepository.org/animsent/vol7/iss32/1/

4. Eisemann, C.H. et al. (1984) ‘Do insects feel pain? – A biological view’, Experientia, 40(2), pp. 164–167. Available at: 10.1007/BF01963580

5. Sneddon, L.U., Elwood, R.W., Adamo, S. and Leach, M. (2014) ‘Defining and assessing animal pain’, Animal Behaviour, 97, pp. 201–212. Available at: 10.1016/j.anbehav.2014.09.007

6. Birch, J, Burn, C, Schnell, A, Browning, H & Crump, A. (2021). ‘Review of the evidence of sentience in cephalopod molluscs and decapod crustaceans.’ LSE Consulting. LSE Enterprise Ltd. The London School of Economics and Political Science. Available at: https://www.lse.ac.uk/News/News-Assets/PDFs/2021/Sentience-in-Cephalopod-Molluscs-and-Decapod-Crustaceans-Final-Report-November-2021.pdf

7. Tracey, W.D., Wilson, R.I., Laurent, G. and Benzer, S. (2003) ‘painless, a Drosophila gene essential for nociception’, Cell, 113(2), pp. 261–273. Available at: 10.1016/S0092-8674(03)00272-1

8. Neely, G.G. et al. (2010) ‘TrpA1 regulates thermal nociception in Drosophila’, Nature, 464(7288), pp. 597–600. Available at: 10.1038/nature08848

9. Hentschel, H. & Penzlin, H. (1982) Beeinflussung des Putzverhaltens bei Periplaneta americana (L.) durch Wundsetzung, Naloxon-, Morphin-, und Met-Enkephalingaben. Zoologische Jahrbücher, Abteilung für Allgemeine Zoologie und Physiologie der Tiere, 86, 535–544.

10. Vierck, C.J., Hansson, P.T. and Yezierski, R.P. (2008) ‘Clinical and pre-clinical pain assessment: are we measuring the same thing?’, Pain, 135(1–2), pp. 7–10. Available at: 10.1016/j.pain.2007.12.008

11. Elwood, R.W. (2019) ’Assessing the potential for pain in crustaceans and other invertebrates’. In: Carere, C. & Mather, J. (eds) The Welfare of Invertebrate Animals. Animal Welfare, vol. 18. Cham: Springer. Available at: 10.1007/978-3-030-13947-6_7

12. Groening, J., Venini, D. & Srinivasan, M.V. (2017) In search of evidence for the experience of pain in honeybees: A self-administration study. Scientific Reports, 7, 45825. 10.1038/srep45825

13. Cammaerts, M.-C. and Cammaerts, R. (2018) ‘Ethological and physiological effects of ibuprofen, the recently most used analgesic: a study on ants as models’, EC Pharmacology and Toxicology, 6(4), pp. 251–267. Available at: https://ecronicon.net/assets/ecpt/pdf/ECPT-06-00159.pdf

14. Fisher, D. N. (2023). ‘Direct and indirect phenotypic effects on sociability indicate potential to evolve.’ Journal of Evolutionary Biology, 36, 209–220. Available at: 10.1111/jeb.14110

15. Association for the Study of Animal Behaviour (ASAB) (2023) Guidelines for the ethical treatment of nonhuman animals in behavioural research and teaching. Animal Behaviour, 195, pp. I–XI. Available at: 10.1016/j.anbehav.2022.09.006

16. Jenkinson, J., Logan, J., Robertson, C., Lockhart, K. and Fisher, D.N. (2026) ‘Behavioural, but not motivational, trade-offs in a cockroach: impact of conditions, injury, and development’, bioRxiv. Available at: 10.1101/2025.10.31.685749

17. Freeberg, T.M., Risner, S.R., Lang, S.Y. and Fiset, S. (2024) ‘Conspecific and heterospecific cueing in shelter choices of *Blaptica dubia* cockroaches’, PeerJ, 12, e16891. Available at: 10.7717/peerj.16891

18. Sakura, M. & Mizunami, M. (2001) Olfactory learning and memory in the cockroach *Periplaneta americana*. Zoological Science, 18(1), 21–28. Available at: 10.2108/zsj.18.21

19. Watanabe, H. and Mizunami, M. (2007) ‘Pavlov’s cockroach: Classical conditioning of salivation in an insect’, PLoS ONE, 2(6), e529. Available at: 10.1371/journal.pone.0000529

20. R Core Team (2024). R: A language and environment for statistical computing. R Foundation for Statistical Computing, Vienna, Austria. Available at: https://www.R-project.org/

21. Kassambara A (2026). rstatix: Pipe-Friendly Framework for Basic Statistical Tests. doi:10.32614/CRAN.package.rstatix. R package version 1.0.0, https://CRAN.R-project.org/package=rstatix

22. Wickham, H. (2016) ggplot2: Elegant Graphics for Data Analysis. New York: Springer-Verlag. Available at: https://ggplot2.tidyverse.org

23. Adamo, S.A. (2019) ‘Is it pain if it does not hurt?’, The Canadian Entomologist, 151(6), pp. 685–688. Available at: 10.4039/tce.2019.49

24. Wada-Katsumata, A., Silverman, J. and Schal, C. (2013) ‘Changes in taste neurons support the emergence of an adaptive glucose-aversion behavior in cockroaches’, Science, 340(6135), pp. 972–975. Available at: 10.1126/science.1234854

